# Unexpected links reflect the noise in networks

**DOI:** 10.1101/000497

**Authors:** Anatoly Yambartsev, Michael Perlin, Yevgeniy Kovchegov, Natalia Shulzhenko, Karina L. Mine, Andrey Morgun

## Abstract

Gene regulatory networks are commonly used for modeling biological processes and revealing underlying molecular mechanisms. The reconstruction of gene regulatory networks from observational data is a challenging task, especially, considering the large number of involved players (e.g. genes) and much fewer biological replicates available for analysis. Herein, we proposed a new statistical method of estimating the number of erroneous edges that strongly enhances the commonly used inference approaches. This method is based on special relationship between correlation and causality, and allows to identify and to remove approximately half of erroneous edges. Using the mathematical model of Bayesian networks and positive correlation inequalities we established a mathematical foundation for our method. Analyzing real biological datasets, we found a strong correlation between the results of our method and the commonly used false discovery rate (FDR) technique. Furthermore, the simulation analysis demonstrates that in large networks, our new method provides a more precise estimation of the proportion of erroneous links than FDR.

## 1. Unexpected correlations

### 1.1. Introducing the concept of unexpected correlations

It is quite common, especially in biology, that in order to understand how system transitions from one state to another (e.g. from health to disease) scientists compare how parameters such as gene expressions, protein levels, or metabolite abundances differ between these states. One result of such a comparison is a list of parameters up- or down- regulated (that is, some numerical value attributed to the parameter has either increased or decreased) from the first state to the second. The parameters are not regulated independently from each other; rather, they make up regulatory networks each with a limited number of key drivers that govern the transition. A common approach to the reconstruction of regulatory network structure is the inference of a correlation network build from these parameters. In particular, correlation (or, for the purposes of this paper, co-variation) networks are widely used in gene expression analysis (see, for example, Butte et al., 2000, Opgen-Rhein and Strimmer, 2007, and references within). Any co-variation network inference implies that any edge in the network (corresponding to correlations between parameter/nodes) is an empirical result of either direct or indirect causal relationships unless they edge is erroneously drawn. The primary question that drove this study was thus whether the causal nature of gene expression networks has any specific implication for their structure and organization. Furthermore, in the case that this relation (causality-network structure) exists, we ask whether it can be used to improve gene network analysis.

In order to address this question we look to basic principles connecting correlation and causality. Causal effects have to follow Reichenbach’s principles (Reichenbach, 1956; Pearl, 2009) which, in the example at hand, imply that if there is a correlation between two genes expressions g_1_ and g_2_, *provided that it is not a statistical artifact*, at least one of three must hold: 1) g_1_ regulates g_2_; 2) g_2_ regulates g_1_; or 3) there is common cause (perhaps another gene, g_3_) that regulates (directly or indirectly) both g_1_ and g_2_ (Figure 1). Thus, in the particular situation under discussion, namely a system with two equilibrium states with two types of regulation (stimulation and inhibition) we propose a scheme in which a sign (positive or negative) of correlation coefficient is associated with direction of regulation of correlated genes. Sign association follows a simple set of rules:

- If there is a correlation between two mutually “up” or “down” regulated genes, the corresponding sign associated with the link is positive.
- If there is a correlation between an “up” regulated gene and a “down” regulated gene, the corresponding sign associated with the link is negative.

**Figure 1:**
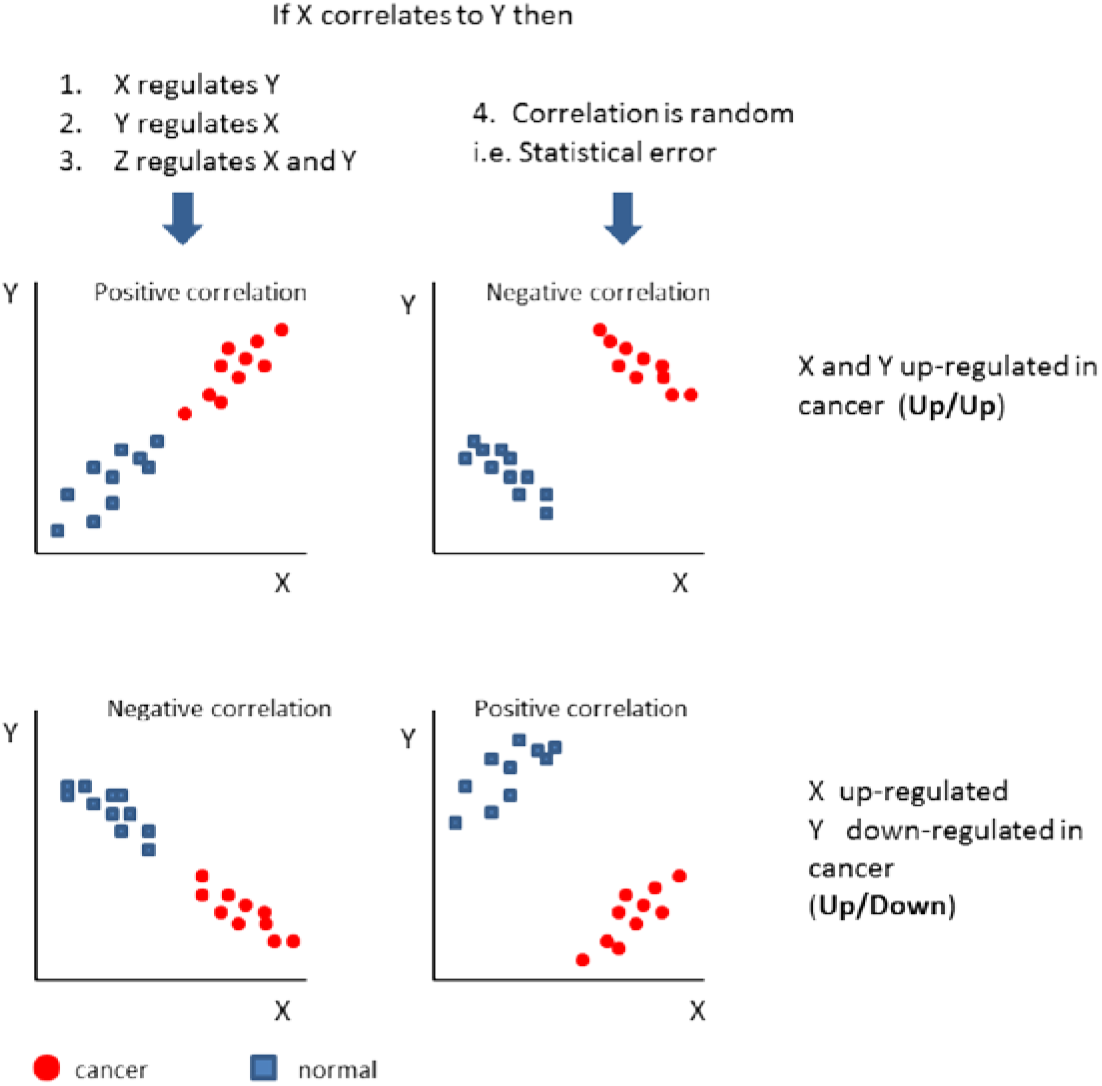
We hypothesize that monotonic regulation mechanisms cannot account for relationships such as the two shown on the right. Correlations exhibiting such regulation between states are thus marked unexpected and hypothesized to result from statistical error.

We hypothesize that correlations whose sign disagrees with that associated with the corresponding link are erroneous (i.e. the result of noise or statistical error rather than causal relationships). We will hereafter call such correlations unexpected, and their rough proportion we abbreviate as PUC (the <u>P</u>roportion of <u>U</u>nexpected <u>C</u>orrelations).

The fundamental reasoning motivating this hypothesis is that regulation mechanisms in biological systems (as well as many other systems) are not generally a function of biological state. Though gene expression levels in a cancerous cell may vary from that in a healthy cell, gene function and regulation schemes in most cases remain constant. The differences in gene expression levels between two biological states should reflect the nature of their regulatory pathways. We expect positively correlated genes to mutually increase or decrease in expression, and negatively correlated ones to be regulated in opposite directions (i.e. up/down or down/up). Note that correlations are evaluated within each state independently, while differences in gene expression is evaluated between the two states. A deviation from this behavior suggests that a particular correlation between the expression levels of two genes is NOT due to a causal link.

A straightforward way to empirically test whether, as we hypothesize, unexpected correlations are erroneous is to analyze some real-world data and compare PUC, which we believe to be a measure of error in a correlation network, to a standard measure of network error, the false discovery rate (FDR) [Benjamini&Hochberg, 1995]. For a proof-of-concept comparison, we used gene expression data from our recently published paper on network analysis in cervical cancer (Mine and Shulzhenko et al., 2013).

We felt that this network should provide excellent real data to analyze our prediction, as it was constructed from a robust meta-analysis of five cancer gene expression datasets and thus validated by large, independent datasets. To our great satisfaction and some surprise, under an FDR threshold of 5% we observed an identical PUC of 5% in this gene expression network (Figure 2, see section I.1. of the supporting material and Figure S1).

**Figure 2:**
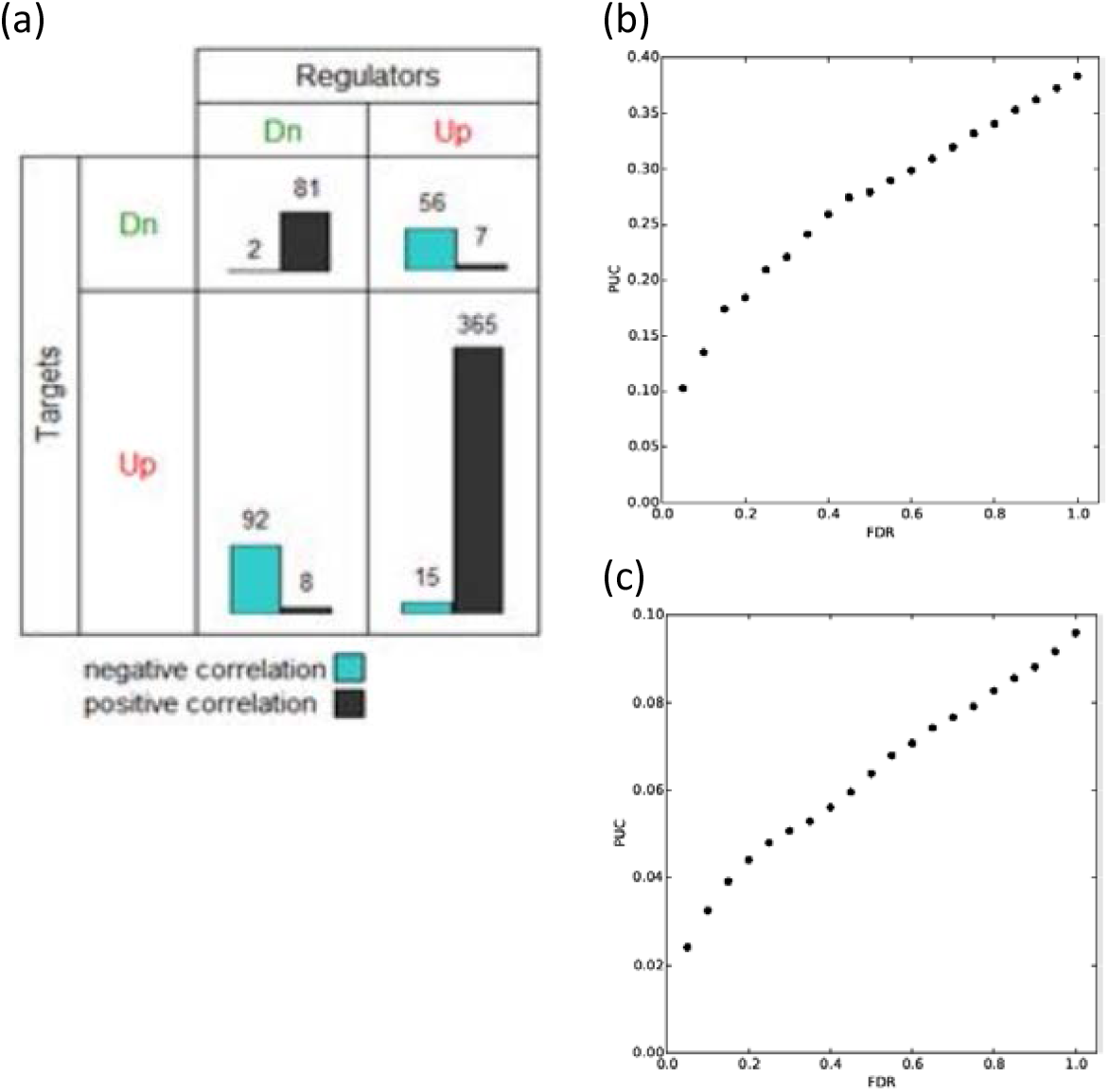
(a) In the initial analysis of cervical cancer gene expression networks, unexpected correlations accounted for 5% of all correlations that passed a 5% FDR threshold. PUC is shown to correlate with FDR in analyzing both gene (b) and macroeconomic (c) data.

The fact that we observed similar levels of unexpected correlations and of erroneous edges in the network reconstructed form cervical cancer data suggests that it can be extrapolated to the whole field of gene-gene regulation and that PUC can potentially be used as a measure of error.

Encouraged by this result, to better understand the properties of this new metric (PUC) we went further to establish a mathematical framework for its application. Indeed, although concept of PUC can be formulated and tested empirically without mathematical theory, a rigorous mathematical formalization of PUC is necessary for its establishment as a widely applicable and powerful method of analysis.

### 1.2. Mathematical formalism relating causation and the sign of correlation

Our hypothesis that unexpected correlations are erroneous can be rigorously proven for systems that transit between two stable states with two types of relations between parameters: stimulation and inhibition. Herein, we provide a proof of our hypothesis in the domain of Bayesian networks (Pearl, 2009) with two equilibrium states and linear dependences between nodes (see proof for more general case in Supplementary Material, section II.2). In order to formulate our results we need to introduce some mathematical notation.

Consider some regulatory network, directed without loops (i.e. a directed acyclic graph, DAG), represented by a graph *G* = (*V, E*). Any edge *e* ∈ *E* is an oriented pair of vertices (nodes) *e* = (*v*, *w*) ∈ *V*^2^. The orientation of an edge represents the direction of causality in a regulatory network (that is, an orientation (*v, w*) implies that *v* regulates *w*). For any node *v* we associate the set of its parents as *pa*(*v*) := {*u* ∈ *V*: (*u, v*) ∈ *E*}. We define the set of grandfathers *gf*(*G*) for the graph *G* as the set of all nodes without parents: *gf*(*G*) := {*v* ∈ *V*: *pa*(*v*) = ∅}.

The graph *G* will be weighted graph. It means that every edge *e* = (*v, w*) ∈ *E* has a label (weight) *c_vw_ ∈* ℝ. With any node *v ∈ V* we associate a random variable *M_v_*. The distribution of random variables is given by their respective structural linear equations *M_v_ = ∑_w_*∈_*pa(v)*_ *c_wv_* *M_w_* + *ε_v_*, where *ε_v_* are mutually independent and identically distributed with mean 0 and variance *σ^2^*.

In the previously discussed biological framework, a graph *G* represents the entire gene expression network. A node *v* represents some gene, which has an expression level *M_v_*. An edge *e* = (*v, w*) represents a causal link between two genes *v* and *w* in which the expression of *w* is regulated by *v*. The sign of *w* reflects the direction of regulation: negative sign and positive sign correspond to inhibition and stimulation, respectively. The parents of *v* are simply all genes which regulate *v* and the grandfathers of *G* are the primary regulators of the entire network, the genes at the top of the regulatory chain.

For simplicity, we consider a regulatory network with only one grandfather (*|gf*(*G*)*| = 1*), denoted by the vertex *o*. 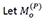 and 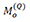 denote the expressions of node *o* in two distinct equilibrium states *P* and *Q*. For any *v* we denote the changes in expression between states as 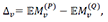, where 𝔼 denotes the expectation value (mean) of corresponding variable.

The mathematical definition of expected and unexpected links, given heuristically in the introduction, is formally expressed in the following way:

#### Definition.

*An edge e ∈ E is called an expected link between nodes v, w ∈ V if and only if* 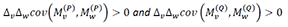. *Any edge which is not an expected link constitutes an unexpected link.*

This definition states that the directions of regulation of two genes between two states should agree with the sign of the correlation between them within each state.

It is straightforward to prove the following lemma (proven in section II.1 of the supporting material):

#### Lemma 1.

*For any finite DAG with linear structural equations there exists some* 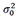 *such that for any variance* 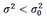 *there are no unexpected links in the graph*.

Lemma 1 implies that in regulatory networks unexpected correlations must have appeared as a result of noise within the network. Thus, the proportion of unexpected correlation thus reflects the noise level in a network.

As a side note, the linear relations between variables can be generalized by the expression 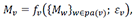 where *f_v_* is some monotonic function over its variables and *ε_v_* is the internal network noise. If the functions *f_v_* are not linear but monotonic, then the lemma still holds.

### 1.3. Unexpected correlations reflect the noise in real and simulated networks.

Mathematical models are restricted by the domain of their assumptions, which may sometimes correspond to only a fraction of real world situations, making them exceedingly limited in applicability. Thus, although we have empirically observed an appropriately small PUC at a low FDR threshold in cervical cancer data, we wanted to verify whether this correspondence would still hold in the gene regulation of an entirely different biological process.

For this we chose a more mundane physiological process than cancer: we analyzed the gene expression network perturbed as a result of colonization of intestinal tissue with normal microbiota (i.e. the mix of microorganisms that live in the gut). In these data, we again found that a low FDR threshold corresponds to a low PUC. Furthermore, PUC is highly correlated with FDR (Figure 2), which provides additional support for our prediction that PUC, similarly to FDR, quantitatively reflects network error.

An important question, however, is whether PUC brings any advantage over the standard approach to measuring the proportion of erroneous edges in a reconstructed regulation network (i.e. FDR). Real data makes such a comparison difficult because though both methods of analysis will return values for network error, there is not necessarily any obvious way to determine which is more accurate; i.e. in real data, the “correct” level of network error is not known.

To investigate the behavior of PUC in a “controlled environment” we simulated Bayesian networks as a model of gene regulation. We define as “true error” any correlation found between the nodes of disjoint, independent networks (Figure 3a).

**Figure 3:**
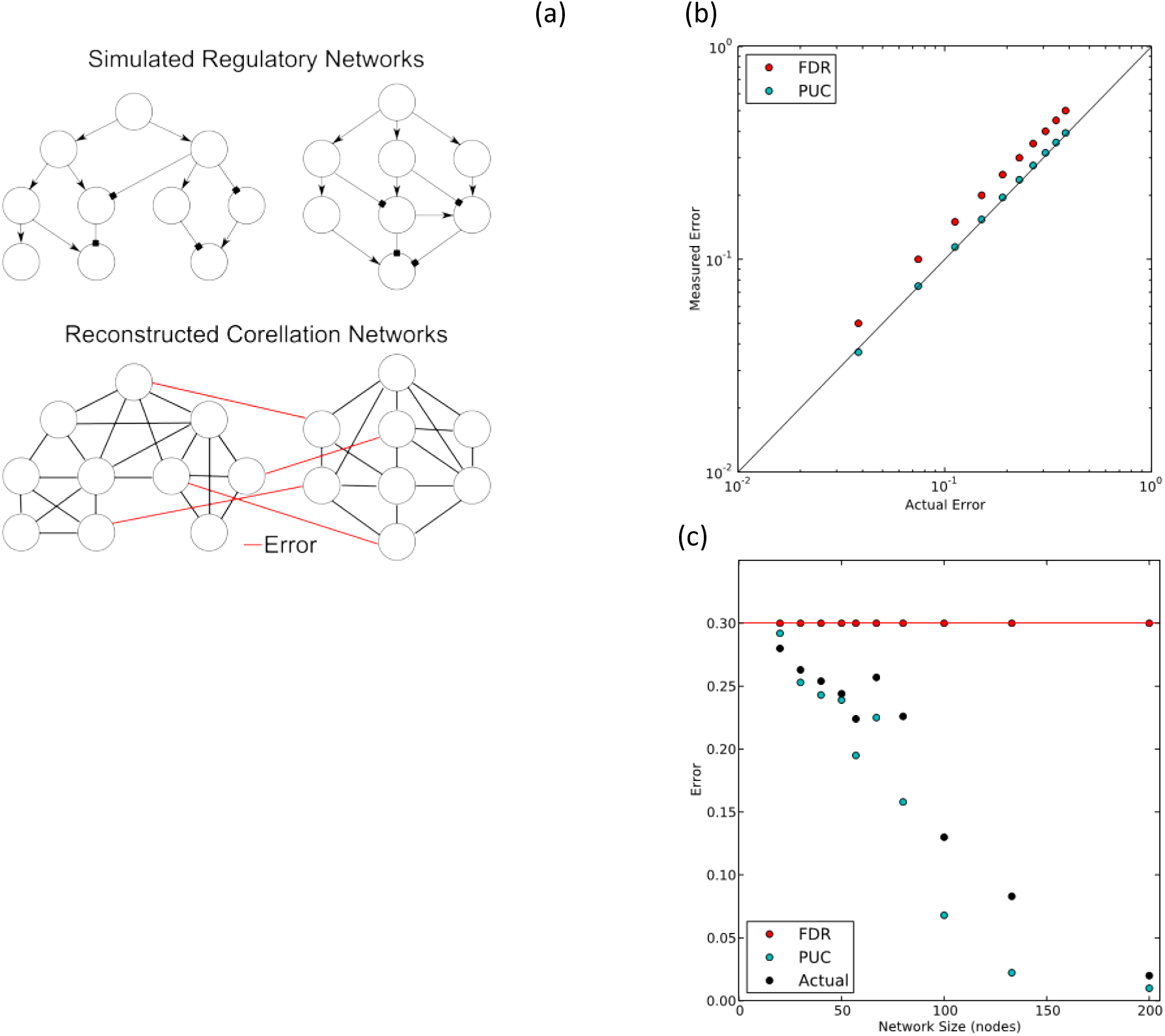
**(a) In order to compare the effectivene of PUC and FDR, two regulatory networks are construct and simulated independently, and both networks’ node expression levels combined into one data set. In reconstructing a correlation network from the simulated data, any correlations between nodes from independent networks are known to be erroneous. This scheme allows for true measure of network error against which to compare PUC and FDR analysis results. (b) Simulations suggest that PUC more accuratel reflects network error than FDR as network size grows, which seems to be due to a more general mathematic feature of PUC (c).**

In order to determine which method (FDR or PUC) better quantifies error, we look at all three measures of error (FDR, PUC, and the true error) and compare the accuracies of FDR and PUC relative to true error (Figure 3b). Simulation results demonstrate that PUC is more accurate than FDR in estimating true error.

It is known that FDR is an overly conservative approach (i.e. it overestimates the number of false positives) in cases when the hypotheses of an analysis are inter-dependent. In the case of regulatory networks, each edge constitutes a hypothesis; interdependency of regulatory network hypotheses manifests in indirect regulation between genes. Indeed, this is exactly the case with co-variation networks, in which it is possible to find numerous indirect pathways with only a few direct links. Using PUC as a measure of error, however, does not require any assumption of hypothesis independence. PUC may thus be more applicable than FDR for reconstruction of networks with a large number of interconnected nodes. The degree of dependency between hypotheses also depends on the size and number of sub-networks that compose a network. A network made up of ten sub-networks consisting of ten nodes each should have a lower degree of hypothesis interdependency than a single network consisting of one hundred nodes lacking any well-defined sub-networks. PUC may thus similarly be more applicable than FDR for analyzing networks with a large edge density. In agreement with these presumptions, we found in simulation analyses that FDR initially provides an accurate estimate of real false positives for small networks (approximately 20-50 nodes, Figure 3c), but diverges from true error as the sizes of networks grow.

We hypothesize that PUC is expected to reflect error independently of size of the network. In order to test this prediction, we performed the same comparisons between the accuracies of FDR and PUC for networks of varying size. The results demonstrated that PUC is more accurate than FDR for larger networks, with differences in accuracy becoming negligible at network sizes of approximately 20 nodes (Figure 3c).

### 1.4. Noise estimation and error correction.

Another very important property of PUC is that it represents approximately half of all erroneous correlations:

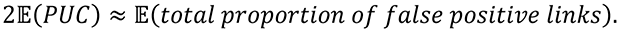

A formal proof of this statement is given in section III.3 of the supporting material, as well as an explanation for why it should make intuitive sense.

The identification of unexpected correlations has two primary impacts. Firstly, it provides a new method to estimate the proportion of erroneous links in a network. Secondly, it allows for the *removal* of approximately half of the erroneous edges in the network (namely, those that are unexpected), decreasing their proportion by a factor of two, thus improving the overall accuracy of the reconstructed network. The final value of network error consists of an estimated proportion of remaining false positive correlations.

The entire procedure for a correlation network is as such: first, all correlations in a differential expression list are ranked by p-value. A network is constructed with edges consisting of correlations within an arbitrary p-value threshold (e.g. 0.01). Unexpected links are identified, counted, and removed from the network. The final error in the remaining network is given by *u/*(*t − u*), where *u* is the number of unexpected correlations and *𝑡* is the total number of correlations within the p-value threshold.

### 1.5. PUC in a non-biological system.

The fact that we could mathematically prove the relationship between unexpected correlations and network error suggests that this principle could be widespread beyond gene interactions in various biological systems. As a proof-of-concept of PUC’s generality, we turned our attention to economics. The basis for this interest was the presumption that economy, similarly to biology, is ruled by cause-effect relationships and, by extension, can be described with regulatory networks. We analyzed 1503 parameters (retrieved from World Bank economic databases) for the year 2008 in 193 countries in such areas as business, education, health, etc. Parameters with bimodal distributions (such as expenditure on primary education as a percent of GDP per capita) defined distinct states of economic networks for any given country. As expected, these networks also demonstrated a high concordance between the network errors given by PUC and FDR (Figure 2C, Figure S2). This result supports the idea that the concept of unexpected correlations can be extrapolated to a large variety of causal networks and that measurement of the proportion of unexpected correlations (PUC) can improve network analysis in many different fields of science.

## Discussion

The growth of molecular biology has advanced such that we can measure the expression of thousands of genes simultaneously. Simply measuring the expression of multiple individual genes, however, is insufficient to describe a systems issue such as complex diseases. To relate gene expression to physiological states (e.g. disease) and other variables in an organism’s environment we utilize gene expression networks. These networks enable more intelligent identification of molecular subtypes of diseases and molecular targets for treatment. The reconstruction of gene expression networks, however, is not easily accomplished. Constructing reliable gene expression networks with current methods requires obtaining large data sets and/or discarding sizeable portions of data to reduce false positive deductions.

Although the False Discovery Rate (FDR - Benjamini-Hochberg, see Benjamini and Hochberg, 1995) is the most popular multiple hypothesis correction procedure, its application for network inference is a conservative procedure and makes the often unfitting assumption of the independence between correlations in gene networks. There are less popular versions of FDR (for example Benjamini-Yukateli) which take into account various dependence structures between the hypotheses under consideration, but the usage of these corrections does not demonstrate any significant advantage over PUC (data not shown). Consequently, these corrections tend to have a rate of high false negative discovery (i.e. low power) and require vast sample sizes in order attain desirable degrees of certainty about reconstructed networks. There is thus a critical need for more powerful methods of estimation of false positive connections between genes in co-expression networks.

In this study we have revealed and mathematically proved a new feature of causal networks. This feature is based on the notion that any correlation has causal and noise components. In the case that causal components prevail over noise, the sign of a correlation between two genes should be related to their up- or down- regulation of the genes between two states (Figure 1). We proposed using this relation for identifying false connections in co-variation networks, increasing network accuracy, an estimating total network error. This approach demonstrates clear advantage over the classic method (FDR) not only by providing better estimates of error in large reconstructed networks, but also by allowing the removal of approximately half of all erroneous edges. The fact that PUC demonstrates similar behavior to standard methods of analysis (i.e. PUC has a strong correlation with FDR) in both real and simulated Bayesian networks further supports the use of this adopted modeling approach. Indeed, certain questions can only be answered using a modeled system. We had to use simulated networks where we know the exact number of false links to compare FDR and PUC.

The concept of expected and unexpected correlations that we introduced is closely related to the concept of monotone causal effects and the covariance between them. The rules we proved for linear relations should therefore hold for any monotone relationships; this idea is expanded in section II.2. of the supporting material, and the framework of PUC extended to a broader class of networks than those mentioned thus far.

We must also address how non-monotonicity affects the notion and application of unexpected correlations. The concept of non-monotonicity can be exemplified for our problem as different types of relationships in two network states, such as a negative correlation between parameters in one biological state and a positive correlation in another. In such cases, despite violation of monotonicity, we expect unexpected correlations to arise primarily due to noise, rather than the change in relationships. Nonetheless, we demonstrated (see section II.4. of the supporting material) that there is no evidence for non-monotonicity to suggest that these exceptionally rare non-erroneous correlations are in fact responsible for the observed changes in gene expression between states of a biological system. Therefore, because the ultimate goal of network inference is actually to model and understand the transition of biological system from one state to another, we can safely remove these unexpected correlations from the reconstructed network for independent reasons (i.e. that they do not have causal contribution to system state transition).

We believe that this work introduces an entirely new way of dealing with error in regulatory network reconstruction. Indeed, statistical methods employed for such problems normally estimate an error, but cannot detect erroneous edges. We propose a method that besides (according to simulations, potentially superior) error estimation allows for identification and removal of approximately half of total network error. Thus, the identification and removal of unexpected correlations decreases the proportion of irrelevant and erroneous connections and strongly increases the power of network inferences.

Finally, our study provides a good example of the success of a systems approach. The collaboration between biologists and mathematicians resulted in the integration of fundamental principles of causality with real world findings (e.g. Figure 2a cervical cancer) to provide the scientific community with a powerful technique that improves the traditional task of network inference from observational data.

## Acknowledgements

We thank Eric Zubriski, Chris Sullivan, and Xiaoxi Dong from Oregon State University for help in setting-up computation infrastructure at CGRB (Center for Genomic Research and Biocomputing).

## Supporting Material

### I. Experimental procedures

#### I.1. Statistically significant correlations between differentially expressed genes (DEGs) show expected signs

In our recent study (Nature Commun. 2013;4:1806) we have shown that key drivers of cervical carcinogenesis are located in regions of frequent chromosomal aberrations and that these genes cause most of the alteration in gene expression in cervical cancer. Therefore, in order to evaluate whether statistically significant correlations between DEGs which result from known causal relations follow our prediction we performed the following analysis:

First, we selected two groups of genes from DEGs discovered in our previous study: 1) genes in which it has been determined that chromosomal aberrations are responsible for the change in regulation; and 2) genes located in regions in which aberrations are rare, defined by FqG–FqL between −0.1 and 0.1 (Figure S1). Next, we analyzed gene co-expression in tumors samples in order to find correlations between those two groups of DEGs. We found 626 correlated gene-gene pairs with FDR 5%. The results provided support to our hypothesis that significant correlations should to have “expected” signs. Indeed, 95% (594 of 626 total pairs) of significant correlations had expected signs.

### II. Theoretical basis.

Here we provide some formal definitions of concepts used in the paper and all necessary proofs. This section consists of four parts: 1) we introduce the mathematical machinery for PUC using Bayesian networks; 2) we generalize the previous formalism to handle a broader set of cases; 3) we demonstrate that PUC reflects half of total network error; and 4) we address concerns with network non-monotonicity.

#### II.1. PUC on Bayesian networks.

In order to apply the new concept of noise estimator we use Bayesian Networks as a convenient model for gene expression. Let *G* = (*V, E*) be some network, which is directed acyclic graph (DAG). Any edge *e ∈ E* is an oriented pair of vertices *e* = (*v, w*): and direction of edge is from the first vertex *v* to the second vertex *w*. We assume that the graph is weighted graph – any edge *e* = (*v, w*) has its labels (weight), *c_vw_*, which is some real number *c_vw_* *∈* ℝ. For any node *v* we associate the set of parents of the node *v*:

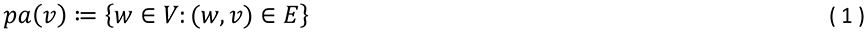

We define the set of grandfathers for the graph *G*:

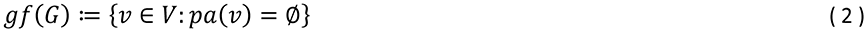

With any node (gene) *v* ∈ *V* we associate the random variable (gene expression) *M_v_*. The random variables satisfy the following linear relations (structure equations): for any *v* ∉ *gf*(*G*)

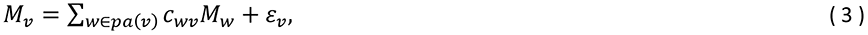

where *ε_v_* are i.i.d. random variable (intrinsic noise) with mean 0 and variance *σ^2^*. Moreover, for simplicity we suppose that there exist only one grandfather *|gf*(*G*)| = *1* and let us denote it as a vertex *o*.

A path *𝜋*(*v, w*) of length *𝑛* from a vertex *v* to a vertex *w* is a sequence of edges *e_𝑖_* = (*v_𝑖_, v_𝑖+1_*), *𝑖* = 1, …, 𝑛 − 1, with *v_0_ = v* and *v_𝑛_ = w*. The weight of the path *𝑊*(𝜋(*v, w*)) is the product of weights of edges from this path:

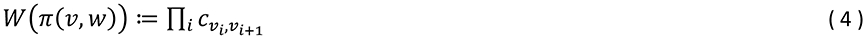

Let *Π*(*v, w*) be the set of all paths connecting nodes *v* and *w*. And let

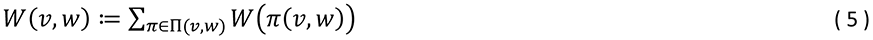

The graph coupled with expressions we consider as a model of regulatory signaling paths system. The distribution of expressions within the system is determined by the topology of graph, weights and the distribution of expressions of grandfathers. Fixed graph and weights the state (joint distribution of variables *M_v_, v ∈ V*) will be defined by the distribution of grandfather.

For example, let *o* be the grandfather vertex and 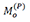 and 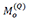 its expressions in these two different states. Same indexes we add for expression for any node in two different states. Denote *𝑑^2^, 𝑑* the variance and standard deviation for grandfather expression in two states, suppose that they do not depend on the state: 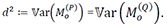 Denote the mean changes in expression of grandfather’s gene as Δ_0_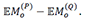. Expression for any non-grandfather vertex *v* can be expressed by formula: for any state *𝑆* ∈ {*P, Q*}

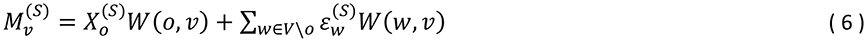

The mean change in expression of gene *v ∈ V\o* is

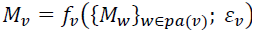

Moreover, for any *𝑆* ∈ {*P, Q*}

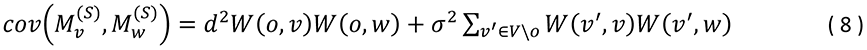

#### Definition.

*We say that a pair of genes v, w ∈ V satisfy **expected correlation inequality** if and only if*

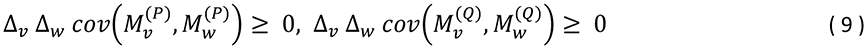

*If (9) holds then we say that the two gene expressions 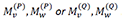 have **expected correlations**. If one or both expected correlations inequalities are not satisfied, we say that 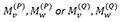 have **unexpected correlations***.

Note that the considered model, by (8) the covariations in (9) do not depend on a state: 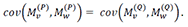. It means that in definition we can use only covariation in one state. In this case the following statement takes place.

##### Lemma 1.

*For any finite DAG network with linear relations between variables there exists some 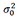 such that for any 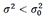 there are no unexpected correlations into the network*.

*Proof*. Direct from formulas (7), (8). By definition (9) and by representations (7), (8) we have

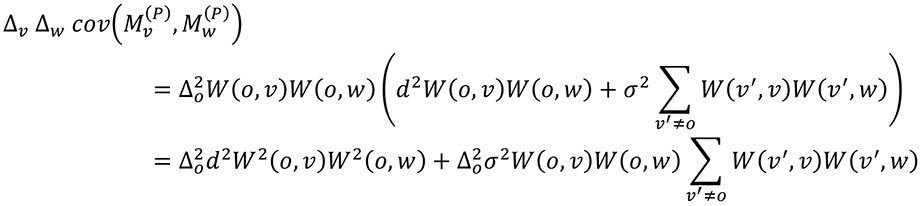

The second sum can be made as less as possible because of *σ^2^*. It proves the Lemma.

The formula (8) shows that any link/correlation between two nodes in a network can be represented as a sum of two parts: *causal propagation* from causal node and *noise propagation* part:

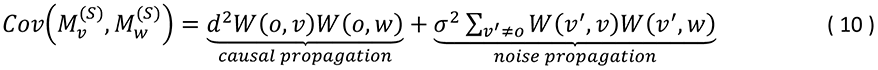

Here, it is easy to see that if the grandfather variance *𝑑^2^* increase, then the causal propagation will determine the sign of the covariance after some threshold. It means that it determines a link to be expected or unexpected.

Moreover, Lemma says that if we observe in such regulation networks (DAGs with linear relationships between variables) unexpected correlations, it means that they appeared as a result of noise propagation within the network. Thus the proportion of unexpected correlation reflects the noise level on a network.

*<u>Note 1.</u> The concept of expected correlations was also observed in VanderWeele and Robins, 2010, as a rule governing the relationship between monotonic links and the sign of covariance between variables*.

*<u>Note 2.</u> The linear relations between variables can be generalized: the expression 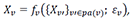, where fv is a monotone function, and ε_v_ is internal network noise. If structural functions are monotonic function, then the lemma holds also*.

*Estimation of noise.* The error estimation based on the following. If two genes belong to two unrelated subnetworks (see Figure 3a), then the correlation between their respective expression levels has to be equal to 0. However, observable correlation can be significantly different from 0 due to noise, in which case, the observable correlation is positive (or negative) in close to 50% of the cases (see formula (20)). Then, on average, half of all random correlations between any pair of genes from unrelated subnetworks can be classified as unexpected, as in (9). Thus *2* *· P𝑈𝐶* can be utilized as an error estimator.

Moreover, it is possible to prove for tree like graphs that within one network the noise propagation (see the formula (11)) has the same property as stated in formula (20). Indeed, the representation (6) means that any variable *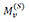* can be decomposed into the causal component *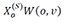* and the noise component *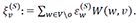*. Then the covariance between 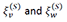 can be calculated exactly (compare with formula (10))

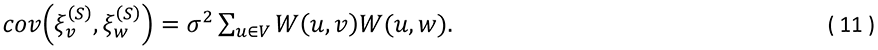

If *c_vw_* are mutually independent, identically distributed, with positive probabilities for being positive or negative, then the covariance (11) for any *𝑆* ∈ {*P, Q*} will be negative approximately in half of cases.

#### II.2. Definitions and generalization.

Here we study the concept of unexpected links in a more general framework. The positive and negative correlation inequalities are an active research direction in the field of probability and statistical mechanics. We believe these inequalities will allow us to generalize the concept of unexpected correlations in the PUC method. The following framework connects FKG (Fortuin–Kasteleyn–Ginibre) inequality in Statistical Mechanics to the concept of expected and unexpected links.

Let Ω be the underlying sample space of a biological system, as an example of a biological system we consider a gene regulatory network, and Ω can be considered as a set of all possible gene expression configurations. We can suppose that the state space Ω has an ordering (or partial ordering) “≺” assigned to pairs of its elements. Here, if *𝜔, 𝜔′, 𝜔″ ∈ Ω*, and if *𝜔 ≺ 𝜔′* and *𝜔′ ≺ 𝜔″*, then *𝜔 ≺ 𝜔″*.

In statistics and in statistical mechanical models the notion of an increasing random variable is remarkable.

##### Definition.

*A random variable 𝑋 = 𝑋*(*𝜔*) *is said to be increasing if 𝜔 ≺ 𝜔′ implies 𝑋(𝜔) ≤ 𝑋(𝜔′). Similarly, a random variable is decreasing if 𝜔 ≺ 𝜔′ implies 𝑋(𝜔) ≥ 𝑋(𝜔′). Both types of random variables, increasing and decreasing, are said to be monotone random variables*.

In the field of statistical mechanics and probabilistic combinatory, the FKG inequality (Fortuin–Kasteleyn–Ginibre inequality) explains most of the results involving monotone random variables and monotone (increasing or decreasing) events. It states that for two increasing random variables *𝑋* and *𝑌*,

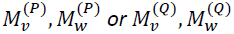

In some applications, such as percolation models, partial ordering of *𝛺* is sufficient for the FKG to hold. See reference [1]. Many important results in applied mathematics and physics, such as the exact value of critical probability in two-dimensional percolation models, would have been impossible without the FKG inequality.

Let *G = (V, E*) be a graph (network) with vertices (nodes) *V* and edges *E*. Nodes *v ∈ V* represent the genes. Let *𝑋_v_(𝜔*) be monotone functions (random variables) assigned to each node *v ∈ V*. Here *𝑋_v_* represents the noiseless gene expressions. In this framework it is convenient represent the state system as a probability measure. Consider two probability measures *P* and *Q* over *𝛺* such that

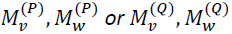

*f* or all *𝜔 ∈ 𝛺*. Here *P* and *Q* correspond to the two states of a biological system. Let us denote, as before, *Δ_v_≔ 𝔼P[𝑋_v_] − 𝔼Q[𝑋_v_]*. Here the variables not have anymore the indices for variables but for expectations with respect to the corresponding measures. We repeat the definition of expected and unexpected links.

#### Definition.

*We say that random variables 𝑋_v_ and 𝑋_𝑢_ modeling gene expressions in a pair of genes satisfy **expected correlation inequality** if and only if*

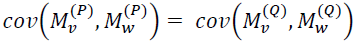

*in which case we say that the two gene expressions 𝑋_v_ and 𝑋_𝑢_ have **expected correlations**. If one or both expected correlations inequalities are not satisfied, we say that 𝑋_v_ and 𝑋_𝑢_ have **unexpected correlations***.

##### Lemma 2.

*If 𝑋_v_ and 𝑋_𝑢_ are monotone functions, and probability measures P and Q satisfy the condition (13), then 𝑋_v_ and 𝑋_𝑢_ satisfy expected correlation inequality (or 𝑋_v_ and 𝑋_𝑢_ have expected correlations*).

*<u>Proof</u>*. Indeed, if *𝑋_v_* is increasing (decreasing) variable, then *Δ_v_≤ 0* (*Δ_v_≥ 0*). Now, if both *𝑋_𝑢_* and *𝑋_v_* are either increasing or decreasing the FKG inequality (12) implies non-negative correlations, so that for any state *𝑆 ∈ {P, Q}*

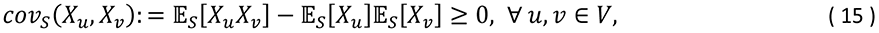

which implies expected correlation inequalities (14).

Similarly, if one of the two variables (i.e. *𝑋_𝑢_* or *𝑋_v_*) is increasing while the other is decreasing, the FKG inequality (12) implies non-positive correlations, such that for any state *𝑆 ∈ {P, Q}*,

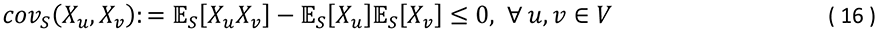

implying (14) hold once again. It proves the Lemma 2.

Next, let *𝜉_v_* denote the errors for each node *v ∈ V*. We assume that the random variables *𝜉_v_, v ∈ V* are functions over a probability space Ξ, independent from any probability measure over *𝛺*, such as *P* and *Q*. Let *𝜇* be the joint distribution of *𝜉_v_, v ∈ V* and 𝔼*_𝜇_[𝜉_v_] = 0* for any *v ∈ V*. The measured gene expression we quantify as a random variable

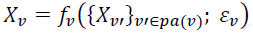

over the product space *𝛺* × Ξ, and the two different states of a biological system correspond to two different probability product measures, *P × 𝜇* and *Q × 𝜇*. Note that for any gene *v*:

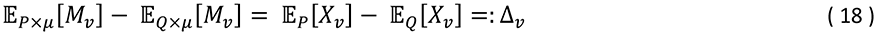

The following Lemma is an analogous of the Lemma 1 for the general framework.

#### Lemma 3.

*If the variances of errors* 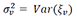 *are small enough for all v ∈ V, then the pairs of measured gene expression M_v_ will also satisfy the inequalities (14). Thus in the noiseless networks we foresee no unexpected correlations*.

*<u>Proof</u>*. The proof is direct consequence of the covariance calculation.

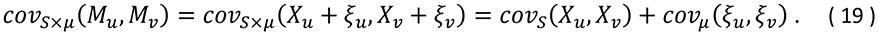

By Cauchy-Schwarz inequality

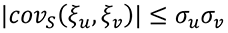

the second covariance in (19) can be made so small that the sign of *cov_𝑆_(M_𝑢_, M_v_*) and the sign of *cov_𝑆_(𝑋_𝑢_, 𝑋_v_*) will coincide. This proves Lemma.

However in the noisy networks, the expected correlations rule (14) can be violated. Here the fraction of edges (*𝑢, v*) violating (14) that we call the Proportion of the Unexpected Correlations (PUC) becomes an estimator of the frequency of false edges.

#### II.3. PUC represents 50% of erroneous.

For any *𝑢, v ∈ V*; *𝑆 ∈ {P, Q};* and *𝜇 ∈* Ξ, let us assume that the variables *𝜉_v_* are random such that, asymptotically, *cov_𝜇_(𝜉_𝑢_, 𝜉_v_*) is positive for half of the 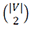 edges (*𝑢, v*), and negative for the rest of the pairs:

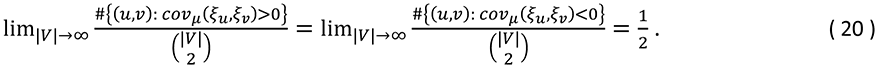

If the covariance *cov_𝑆×𝜇_(M_𝑢_, M_v_*) is of a different sign than *cov_𝑆_(𝑋_𝑢_, 𝑋_v_*) (i.e. if a particular correlation (*𝑢, v*) is unexpected), it must hold that (see (20)):

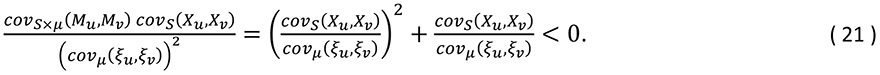

This condition is of the form *𝑅^2^ + 𝑅 < 0*, where 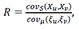 which trivially has the solution:

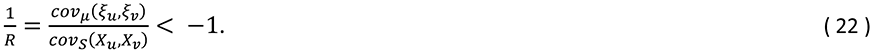

The resulting inequality is satisfied under two conditions, which are thus requisite for a correlation to be unexpected, namely:

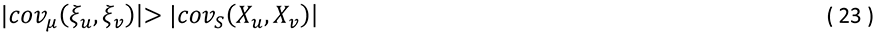

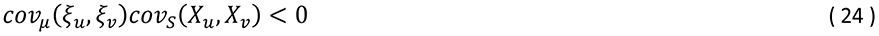

The first condition (23) is interpreted as a drowning out of the causal link between two nodes by error; that is, the magnitude of error in the correlation between two nodes’ expressions is greater than the magnitude of real correlation between them. The second condition (24) is interpreted as a counteracting of error to causal connections: the contribution to the empirical correlation between two nodes due to error must counteract the contribution due to causal mechanisms.

Condition (24) implies that, given a condition (20) for error distribution, PUC will statistically detect 50% of total false correlations for which the causal contribution is negligibly small, as the signs of the error and causal contribution are equally likely to be the same as they are to be opposite.

#### II.4. Unexpected correlations under non-monotonicity.

Here we prove the proposition in the conclusion about non-monotonic links. The statement says that a non-monotonic link between two nodes with an unexpected correlation cannot cause a transition between two distinct states of a network. We provide an extreme example of non-monotonicity, in which the dependence between two nodes changes in sign in the two states of a network (e.g. stimulation in one state of a biological system and inhibition in the other).

Assume we are given *𝑛 +* 2 *gene expressions in two biological state P* and *Q*: *𝑋_P_, 𝑌_P_, 𝑋_1,P_, …, 𝑋_𝑛,P_* and *𝑋_Q_, 𝑌_Q_, 𝑋_1,Q_, …, 𝑋_𝑛,Q_*. We assume linear (or almost linear) dependence of *𝑌* on *𝑋* within any one given biological state, stated as follows: *𝑌_P_ = 𝛼_P_𝑋_P_ + 𝜉_P_* and *𝑌_Q_ = 𝛼_Q_𝑋_Q_ + 𝜉_Q_*, where *𝜉_P_* is a function of *𝑋_1,P_, …, 𝑋_𝑛_,P*, and *𝜉_Q_* is a function of *𝑋_1,Q_, …, 𝑋_𝑛,Q_*, and *𝛼_P_𝛼_Q_ ≠ 0*. We suppose that *𝑋_P_* (*𝑋_Q_*) and *𝜉_P_* (*𝜉Q*) are independent. Recall that all gene expression values are positive and remember that *Δ𝑋* ≔ 𝔼*_P_*[*𝑋*] − 𝔼*_Q_*[*𝑋*] = 𝔼[*𝑋*_*P*_] − 𝔼[*𝑋_Q_*].

##### Lemma 4.

*Suppose 𝛼_P_𝛼_Q_* < 0 (*implying that the relation between X and Y is non-monotonic), then:*

a. *𝑋 and 𝑌 have unexpected correlations*.
b. *The sign of Δ𝑌 may not depend on the sign of Δ𝑋, but instead mostly depends on the sign of Δ𝜉*.

*<u>Proof</u>*. Observe that, due to independence of *𝑋_P_* (*𝑋_Q_*) and *𝜉_P_* (*𝜉_Q_*):

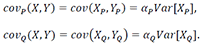

Therefore, *cov_P_*(*X, Y*)*cov_Q_*(*X, Y*) < 0 (*so that the expected correlation inequalities do not hold simultaneously) if and only if* *𝛼_P_𝛼_Q_* < 0. This proves the item (a) of the lemma.

Let us prove (b). Without loss of generality, *cov_P_(𝑋, 𝑌)* < 0, *implying* *𝛼_P_* < 0 *and* *𝛼_Q_* > *0*. Hence:

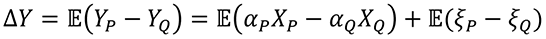

Note that 𝔼(*𝛼_P_𝑋_P_ − 𝛼_Q_𝑋_Q_*) *<* 0 *regardless of the values of* *𝑋_P_* and *𝑋_Q_* (both of which are strictly positive). Thus in the case *Δ𝜉 > 0* the change *Δ𝑌* will still be negative. The sign of *Δ𝑌* will be positive only if *Δ𝜉 ≫* 0.

---

**Figure S1:**
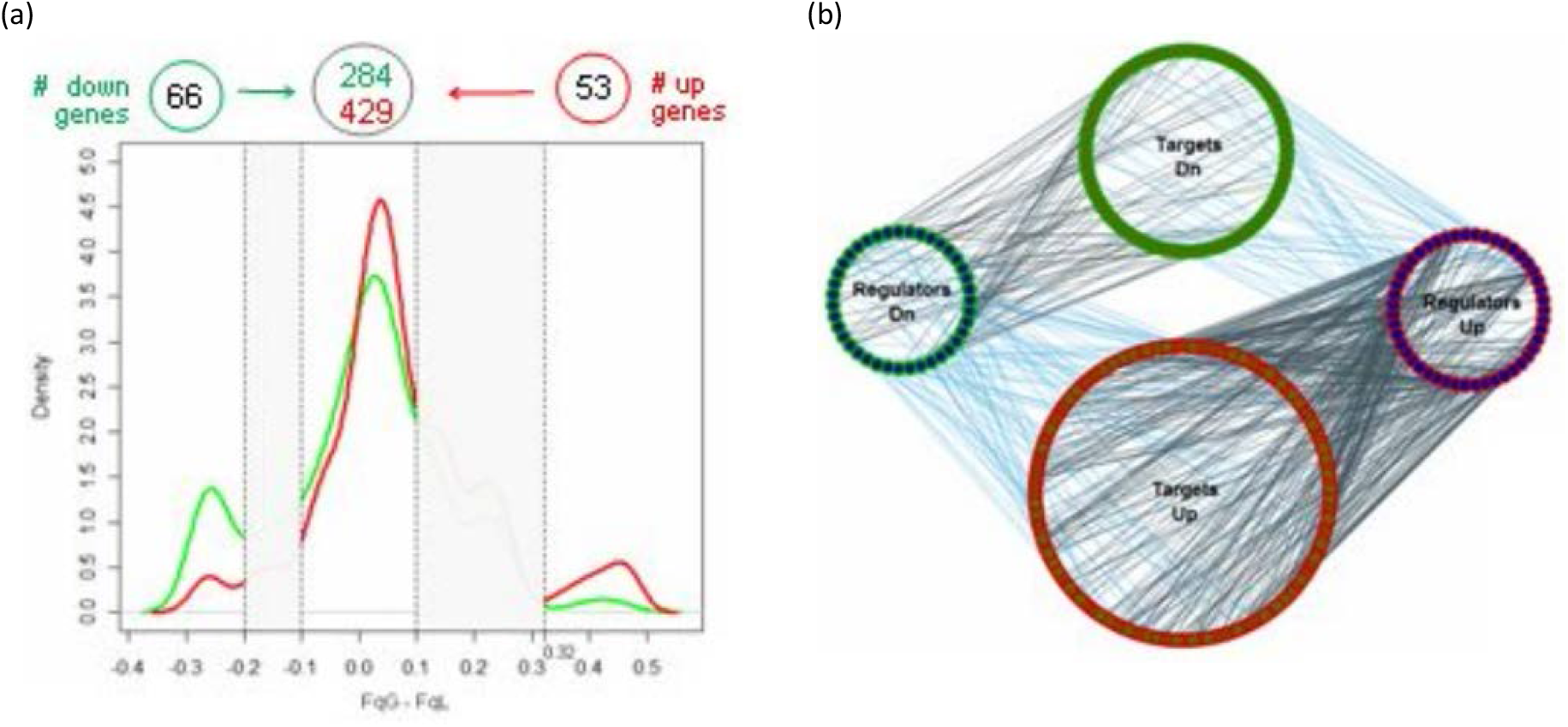
Genes directly regulated by chromosomal aberrations can also in turn regulate genes located outside of the aberrations. (a) Genes regulated by chromosomal aberrations in the expected direction (located in the regions *FqG − FqL < −* 0.2 *or* *FqG − FqL* > 0.3) *were considered as potential regulators, and genes located within the regions of very rare aberrations* (|*FqG - FqL*| ≤ 0.1) *were considered to be potential targets. The green (red) line represents up-regulated (down-regulated) genes. (b) The reconstructed regulatory network with correlations in agreement with gene expression. The two green (red/purple) circles are made of up down-regulated (up-regulated) nodes, the middle (side) circles are made up of targets (regulators), and the black (cyan) lines represent positive (negative) correlations*.

**Figure S2:**
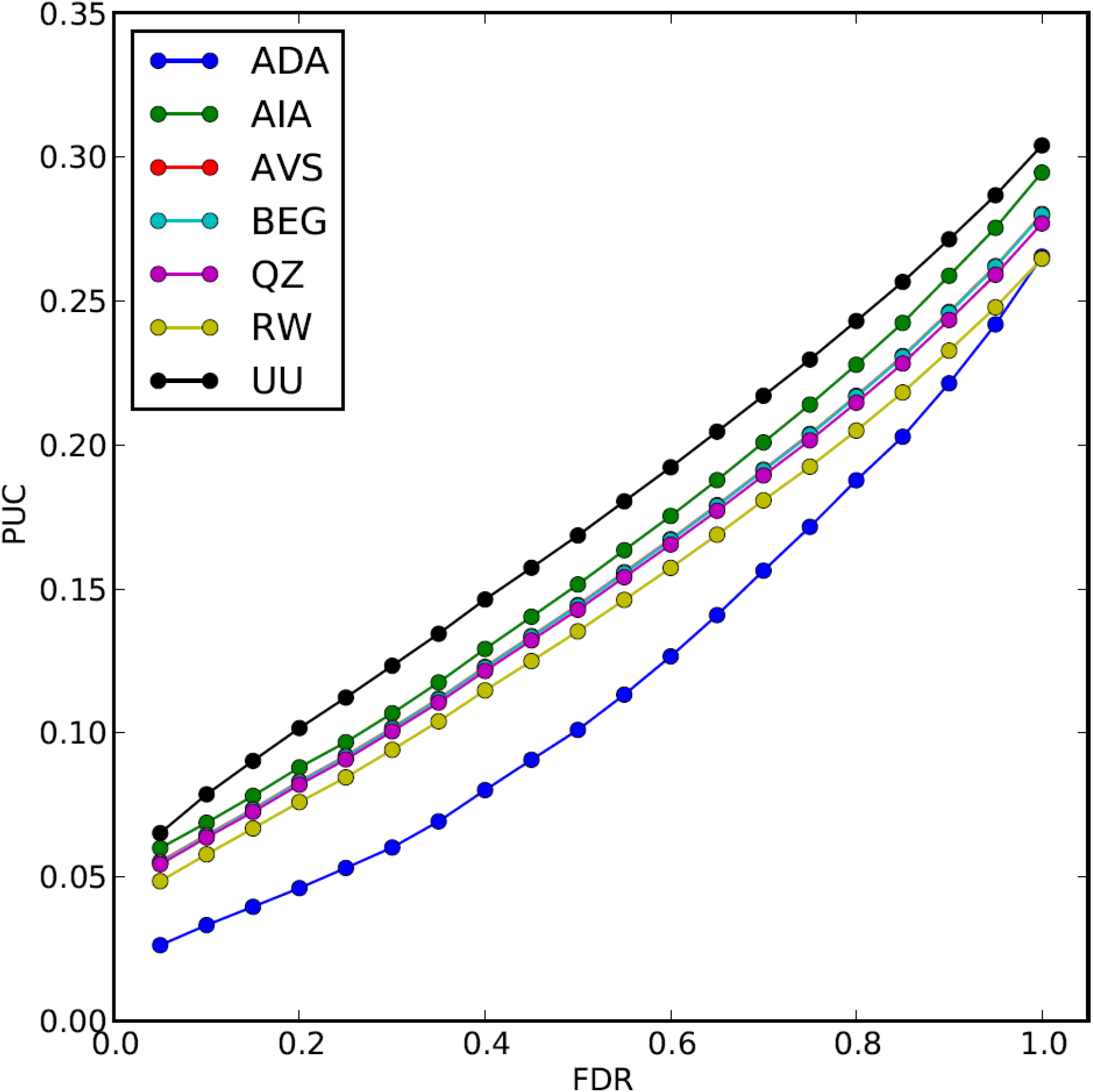
PUC and FDR correlate strongly when reconstructing macroeconomic networks using various bimodal parameters to define system states. Parameters shown are: ADA - Duration of compulsory education; AIA - Cause of death, by communicable diseases and maternal, prenatal and nutrition conditions (% of total); AVS - Manufactures exports (% of merchandise exports); BEG - Educational expenditure in pre-primary as % of total educational expenditure; QZ - Private credit bureau coverage (% of adults); RW - Strength of legal rights index; UU Passenger cars (per 1,000 people)

